# Explaining an early ground-nesting bee emergence with climate and occurrence data: Temperature, not precipitation, advances onset

**DOI:** 10.64898/2026.11.23.768685

**Authors:** Charles N. Thrift, Madeleine M. Ostwald, Jennifer Schlauch Saiyawong, Katja C. Seltmann

## Abstract

Understanding the drivers of bee phenology is important for predicting how climate change may alter the temporal overlap between bee activity and plant flowering, with consequences for pollination. We documented unusually early emergence of *Melissodes tepidus timberlakei* Cockerell, 1926 at a perennial nesting site in spring 2026 and examined relationships between emergence and climatic conditions with (1) five years of nesting observations at a single site, and (2) 119 years of *M. tepidus* occurrence records from across the western United States. We found that warming temperatures were associated with earlier emergence, whereas precipitation was not, and we emphasize the utility of monitoring ground nests to understand emergence drivers in bees during ongoing climate change.

## Introduction

Understanding the drivers of emergence and flight timing in bees (Hymenoptera: Anthophila) is crucial to the conservation of bee–plant interactions amid ongoing climate change. Of the roughly 25,000 bee species estimated worldwide (Dorey *et al*., 2026), most are solitary and univoltine, with adults active for only a few weeks following emergence (Danforth *et al*., 2019). Even small shifts in emergence timing can alter adult flight periods, disrupting synchrony with host plant flowering, resulting in phenological mismatch (Freimuth *et al*., 2022). Such mismatches can disrupt pollination and negatively affect both bee and plant reproduction (Kudo & Ida, 2013; Kudo & Cooper, 2019; de Manincor *et al*., 2023; Schenk *et al*., 2018), potentially increasing extinction risk (Peng *et al*., 2025). Global climate change continues to increase the frequency and intensity of extreme heat and precipitation events (Seneviratne *et al*., 2021), with largely unknown consequences for bee and plant phenology.

Emergence timing can be cued by environmental conditions during development, but in the limited studies conducted to date, bee species do not respond uniformly. Only 113 species from 20 genera have been studied for relationships between emergence timing and temperature or precipitation (Table 1), and responses vary. Other studies include additional bee species, but species-level emergence–climate relationships cannot be determined from studies that analyze shifts in phenology over time rather than with environmental cues (*e*.*g*., Dorian *et al*., 2023) or analyze emergence–climate trends at a community level (*e*.*g*., Stemkovski *et al*., 2020). Within species, interannual variation in emergence timing has been linked to environmental conditions through experimental (Danforth, 1999; Fründ *et al*., 2013; Kemp & Bosch, 2005; Slominski & Burkle, 2019; Vinchesi *et al*., 2013; White *et al*., 2009) or observational studies, using multiple years of natural history and climate data (Bartomeus *et al*., 2011; Olliff-Yang & Mesler, 2018; Robbirt *et al*., 2014; Wyver *et al*., 2023). Overall, most species studied exhibit earlier emergence under warmer conditions, although some exhibit delayed emergence, no detectable response, or conflicting results between studies. In a community-level analysis of 67 bee species in a subalpine environment, emergence advanced with warming temperature and delayed with increased summer rainfall (Stemkovski *et al*., 2020). Precipitation is generally less explanatory for emergence time than temperature (Olliff-Yang & Mesler, 2018), except in some desert bees (Danforth, 1999; Rust, 1988). Species-specific variation and the limited phylogenetic breadth of bees sampled hinder our ability to predict how most bees will respond phenologically to ongoing climate change.

**Table 1.** Bee emergence time response to increased temperature (as ambient air temperature unless otherwise noted) or precipitation in 113 species from 20 genera in 10 studies. “Advance” means emergence became earlier with increased temperature, “delay” means emergence became later with increased temperature, and “no effect” means no significant relationship was found.

| family | genus | species | temp. | precip. | citation |
|---|---|---|---|---|---|
| Andrenidae | *Andrena* | *Andrena barbilibris, A. bicolor, A. chrysosceles, A. carlini, A. cineraria, A. clarkella, A. crataegi, A. denticulata, A. dorsata, A. flavipes, A. florea, A. fucata, A. fulva, A. fuscipes, A. haemorrhoa, A. helvola, A. humilis, A. labialis, A. labiata, A. miserabilis, A. minutula, A. nigroaenea, A. nitida, A. ovatula, A. praecox, A. scotica, A. subopaca, A. synadelpha, A. thoracica, A. trimmerana* | advance |  | 1, 6, 10 |
|  | *Perdita* | *Perdita portalis* |  | advance | 2 |
| Apidae | *Anthophora* | *Anthophora bimaculata, A. plumipes* | advance |  | 10 |
|  | *Bombus* | *Bombus bimaculatus, B. bohemicus, B. campestris, B. hortorum, B. impatiens, B. jonellus, B. lapidarius, B. monticola, B. pascuorum, B. pratorum, B. rupestris, B. sylvestris* | advance |  | 1, 10 |
|  |  | *B. distinguendus, B. humilis, B. ruderarius* | no effect |  | 10 |
|  | *Epeolus* | *Epeolus cruciger* | no effect |  | 10 |
|  |  | *E. variegatus* | advance |  | 10 |
|  | *Habropoda* | *Habropoda miserabilis* (soil temperature) | advance | advance | 5 |
| Colletidae | *Colletes* | *Colletes inaequalis* | advance |  | 1 |
|  | *Hylaeus* | *Hylaeus brevicornis* | no effect |  | 10 |
|  |  | *H. communis* | advance or delay |  | 3, 10 |
|  |  | *H. confusus, H. dilatatus, H. hyalinatus* | advance |  | 10 |
| Halictidae | *Halictus* | *Halictus rubicundus, H. tumulorum* | advance |  | 10 |
|  | *Lasioglossum* | *Lasioglossum albipes, L. calceatum, L. fulvicorne, L. lativentre, L. leucopus, L. leucozonium, L. malachurum, L. minutissimum, L. morio, L. parvulum, L. pauxillum, L. smeathmanellum, L. villosulum, L. zonulum* | advance |  | 10 |
|  |  | *L. laevigatum* | no effect |  | 10 |
|  | *Nomia* | *Nomia melanderi* (soil temperature) | advance |  | 8 |
|  | *Sphecodes* | *Sphecodes crassus, S. ephippius, S. geoffrellus, S. gibbus, S. hyalinatus, S. monilicornis, S. pellucidus, S. puncticeps* | advance |  | 10 |
| Megachilidae | *Anthidium* | *Anthidium manicatum* | advance |  | 10 |
|  | *Chelostoma* | *Chelostoma florisomne* | advance |  | 3 |
|  | *Coelioxys* | *Coelioxys mandibularis* | delay |  | 3 |
|  | *Heriades* | *Heriades truncorum* | delay |  | 3 |
|  | *Megachile* | *Megachile centuncularis, M. ligniseca; M. versicolor* | advance or delay |  | 3, 10 |
|  |  | *M. lapponica, M. maritima, M. relativa, M. willughbiella* | advance |  | 7, 10 |
|  |  | *M. leachella* | no effect |  | 10 |
|  | *Osmia* | *Osmia atriventris, O. aurulenta, O. bicolor, O. bicornis, O. bucephala, O. cornifrons, O. cornuta, O. pumila* | advance |  | 1, 3, 9, 10 |
|  |  | *O. caerulescens, O. californica, O. coloradensis, O. montana, O. tersula, O. vallicola* | no effect |  | 7, 10 |
|  |  | *O. lignaria* | advance or no effect |  | 1, 4, 7 |
| Melittidae | *Dasypoda* | *Dasypoda hirtipes* | advance |  | 10 |
|  | *Melitta* | *Melitta leporina, M. tricincta* | advance |  | 10 |

**Footnote**. Citations are as follows: (1) Bartomeus *et al.,* 2011. (2) Danforth, 1999. (3) Fründ *et al.,* 2013. (4) Kemp & Bosch, 2005. (5) Olliff-Yang & Mesler, 2018. (6) Robbirt *et al.,* 2014. (7) Slominski & Burkle, 2019. (8) Vinchesi *et al.,* 2013. (9) White *et al.,* 2009. (10) Wyver *et al.,* 2023.

A challenge in understanding relationships between emergence and environmental conditions is the inherent difficulty of estimating phenological onset from occurrence data (Park *et al*., 2025). Occurrence data document the presence of a taxon at a location and time based on observations or collected specimens (Wieczorek *et al*., 2012). Perennial ground-nesting sites offer a solution: emergence dates can be recorded based on the appearance of nest holes, and monitoring the same site over multiple years controls for site-specific variation. Notable shifts in emergence timing, especially at well-studied sites, provide opportunities to investigate the environmental factors influencing bee emergence.

Here, we document unusually early adult activity of *Melissodes tepidus timberlakei* Cockerell, 1926 (Apidae) at a perennial nesting site and investigate whether temperature and precipitation explain variation in emergence timing. First, we compared the earliest recorded adult activity from 2022 to 2026 with temperature and precipitation during the presumed developmental period, relative to 30-year climate normals. Based on these observations, we hypothesized that temperature would be a stronger predictor of emergence timing than precipitation. Second, we tested this hypothesis using *M. tepidus* occurrence records from across the western United States (1907–2026), using the earliest collection date for each locality–year combination as a proxy for emergence, alongside climate data from the preceding developmental period (July–May).

## Material and Methods

*Melissodes tepidus* Cresson, 1878 is a univoltine, solitary, polylectic (dietary generalist), ground-nesting bee with three currently recognized subspecies: *M. tepidus timberlakei, M. tepida tepida*, and *M. tepida yumensis* (LaBerge, 1956). A perennial nesting aggregation of *M. tepidus timberlakei* occurs on the shoreline of a coastal lagoon at the University of California, Santa Barbara campus (34.4106 N, 119.8505 W). This nesting site was first described by Ostwald *et al*. (2024) (Figs. 1–3) and has been observed *in situ* since 2016 (Seltmann, pers. obs.). Each year, several hundred nests occur within 2 m of the water on flat, bare soil and among the salt marsh vegetation, which is dominated by saltgrass (*Distichlis spicata*), alkali heath (*Frankenia salina*), Pacific glasswort (*Salicornia pacifica*), and marsh jaumea (*Jaumea carnosa*). The soil at this site is classified as moderately eroded 85% Concepcion fine sandy loam (Walkinshaw *et al*., 2023). Adult bee activity at this site is most common in June and July, consistent with previously documented activity periods. LaBerge (1956) described *M. tepidus timberlakei* as active from April (“presumably the latter part of the month,” LaBerge, 1956, p. 1127) to October 23, with main activity in June and July. The other two subspecies exhibit similar activity patterns. *M. tepida tepida* is active May 26–August 25, with peak activity in July, and *M. tepida yumensis* is active May 12–October 19, with peak activity in June and July (LaBerge, 1956).

**Fig. 1–3:**
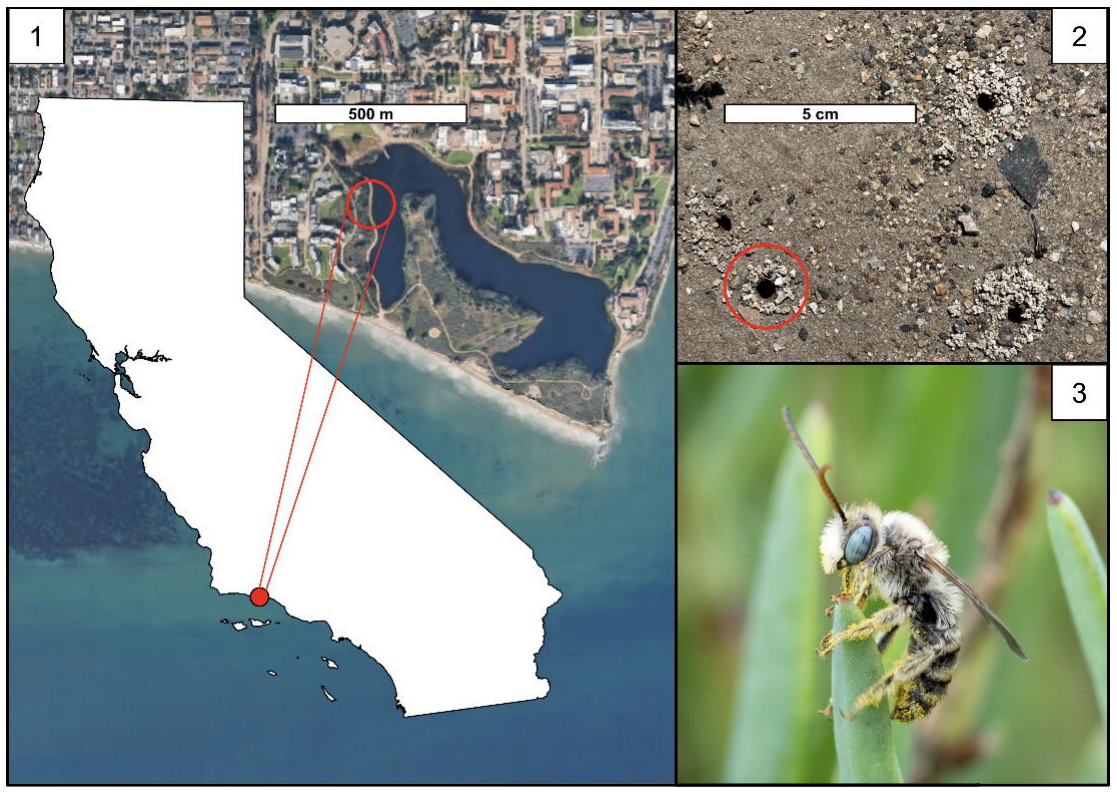
Nesting aggregation site at University of California, Santa Barbara Campus Lagoon (34.4106, –119.8505) (**Fig. 1**), visible nest holes (one circled in red) with 5 cm scale (**Fig. 2**), and *M. tepidus timberlakei* (male) photographed by Jeremiah Bender (**Fig. 3**).

We used two complementary analyses to examine relationships between emergence timing and climatic conditions: observations from a perennial nesting site and historical occurrence records. All analyses were conducted in R version 4.6.1 (R Core Team, 2026). Code and data are available on Zenodo: http://www.doi.org/10.5281/zenodo.23985572

### Emergence with five years of nesting activity (five-year emergence)

Nest-site activity was documented between 2022 and 2026 through field observations and specimen collection intended to capture the full emergence period. We used these records to determine the earliest documented date of adult activity each year. Collected specimens are housed in the University of California, Santa Barbara Invertebrate Zoology Collection at the Cheadle Center for Biodiversity and Ecological Restoration. Specimen records are databased and publicly available through Bee Library (Seltmann *et al*., 2021) and GBIF (Cheadle Center, 2026). We used PRISM temperature and precipitation data to characterize climatic conditions at the nesting site during the presumed developmental period, defined as July of the preceding year through the date of first recorded adult activity in the following spring (PRISM Group, 2026).

We selected July as the starting point to standardize comparisons across years because adult bees were observed provisioning nests during this month in all years, although provisioning activity also occurred during other months depending on emergence timing. We extracted daily maximum temperature (Tmax) and precipitation data for the nesting site from PRISM for 28 June 1991–23 April 2026. We calculated seven-day rolling averages from the daily values for 1 July 1991–20 April 2026. For each day, the seven-day rolling average was calculated as the mean of the values for that day, the three preceding days, and the three following days. Seven-day rolling averages are commonly used in climate research to characterize sustained thermal conditions, including heatwaves, while minimizing variation driven by single-day weather events (Cuffney & Brightbill, 2008). For each developmental period, we calculated cumulative precipitation as the sum of daily precipitation since 1 July. This approach aligns with the developmental period of *M. tepidus* and captures interannual variation in precipitation under the region’s Mediterranean climate, which is characterized by wet winters and dry summers (Seager *et al*., 2019).

To quantify anomalous weather, we compared the observed weather (July 2021–April 2026) to historical 30-year climate normals (July 1991–June 2021). For each day, we calculated the percentile of the seven-day rolling average Tmax and cumulative precipitation relative to the corresponding values in the climate normals. Higher percentiles indicate that an observed value exceeds a greater proportion of the historical values for the same calendar day. A percentile of 100 indicates that the observed value exceeded all 30 historical values, whereas a percentile of 50 indicates that it exceeded half of the historical values. We defined heatwaves as periods when the seven-day rolling average Tmax exceeded the 90th percentile of historical climate normals for five or more consecutive days. The 90th-percentile threshold identifies extreme heat (Ghatak *et al*., 2017; Perkins & Alexander, 2013), whereas the five-day minimum duration distinguishes sustained heat events from short-term temperature spikes (Padalino *et al*., 2024). Similarly, we defined extreme precipitation periods as periods when cumulative precipitation exceeded the 90th percentile of historical cumulative precipitation values for five or more consecutive days. For each of the five emergence years, we counted the number of hot days, heatwaves, high-precipitation days, and extreme-precipitation periods.

### Estimated emergence with 119 years of occurrence data (occurrence data)

We downloaded *M. tepidus* occurrence records from Bee Library (Bee Library, 2026) and a global, validated bee occurrence dataset (Dorey *et al*., 2023). To maximize sample size, we included records identified as *M. tepidus* or any of its three subspecies. We combined records from both sources, removed duplicates, and excluded records lacking a collection date or location, leaving 998 records. We then excluded records from outside the western United States, defined here as California, Arizona, Utah, Nevada, Idaho, and Oregon, leaving 972 records. We removed records that lacked a specified collection month or day, as well as records with collection dates on the first day of a month (n = 57), following Belitz *et al*. (2023), because records with incomplete collection dates may be assigned the first day of the collection month. For example, a collection date of April 2001 but no specified day could be assigned a date of 1 April 2001. Next, we rounded latitude and longitude to two decimal places and removed duplicate records collected on the same date at the same location. Rounding latitude and longitude allowed us to identify geographically close occurrences (within approximately 1.1 km) that did not have identical recorded coordinates. Because climate data were available at 800-m resolution, this step also reduced the likelihood that the same climate data would be assigned to multiple occurrence record locations. This resulted in 381 records. When multiple occurrences were recorded at the same location in the same year, we retained only the earliest collection date and removed others, resulting in 248 records. Retaining the earliest collection date provides a closer approximation of emergence because collection dates may fall anywhere within the adult flight period. We removed the single record collected before 1895, the earliest year PRISM climate data are available. The final dataset comprised 247 occurrence records housed in 14 institutions (Table 2). This exceeds a sample size threshold (n = 100) previously used to quantify phenological shifts in solitary bees using occurrence data (Dorian *et al*., 2023).

**Table 2.** Institutions housing *M. tepidus* records used in this study.

|  |
|---|
| American Museum of Natural History (AMNH) |
| C.A. Triplehorn Insect Collection, Ohio State University, Columbus, OH (OSUC) |
| Cornell University Insect Collection (CUIC) |
| Essig Museum of Entomology, University of California Berkeley (EMEC-UTB) |
| Illinois Natural History Survey (INHS) |
| University of Kansas Natural History Museum Entomology Division (KU) |
| University of Arizona Insect Collection (UA) |
| University of California Santa Barbara Invertebrate Zoology Collection (UCSB-IZC) |
| UCSD Holway Lab - Longterm Habitat Fragmentation Study (UCSD), currently housed at UCSB-IZC |
| University of New Mexico Museum of Southwestern Biology (UNM) |
| United States Department of Agriculture Agricultural Research Service U.S. National Pollinating Insects Collection (USDA-ARS) |
| USGS Native Bee Inventory and Monitoring Lab (USGS PWRC) |
| Smithsonian Institution, National Museum of Natural History (USNM) |
| J.B. Wallis / R.E. Roughley Museum of Entomology (WRME) |

We used PRISM to assign mean temperature, maximum temperature, and precipitation values to each occurrence record. In contrast to our five-year emergence analyses, we used monthly rather than daily climate data because PRISM daily data are unavailable before 1981, whereas our records extend back to 1907. We also used monthly values for records dating after 1981, when daily PRISM data were available, to maintain consistency across the dataset. For each occurrence, we calculated mean temperature over four periods: the developmental period (July–May), fall (September–November), winter (December–February), and spring (March– May). We used May as a standardized endpoint for the developmental period because emergence dates varied among records. For each period, we also calculated mean maximum temperature (the average of monthly maximum temperatures) and cumulative precipitation. Because temperature variables (mean and mean maximum temperature across seasons) were correlated with one another, as were precipitation variables, we retained only two predictors for analysis: mean temperature and cumulative precipitation during the developmental period (July–May).

Using these occurrence and climate data, we tested for relationships between emergence date and climate in *M. tepidus* across the western United States (n = 247). We fitted two linear mixed-effects models (LMMs) using the “lme4” package (Bates *et al*., 2015). In each model, emergence date (Julian day, 1–366) was the response variable, and one climate variable (mean temperature or cumulative precipitation during the developmental period) was included as a fixed effect, with location as a random effect. We assessed model residuals using the “DHARMa” package (Hartig, 2016). We obtained p-values for the fixed effects using the “lmerTest” package (Kuznetsova *et al*., 2017), which approximates denominator degrees of freedom using the Satterthwaite method (Satterthwaite, 1946). We applied a Bonferroni correction by multiplying each p-value by the number of tests (two) and comparing the adjusted p-values with a significance threshold of α = 0.05. We quantified model explanatory power using marginal (R^2^_m_) and conditional (R^2^_c_) coefficients of determination calculated with the “performance” package (Lüdecke *et al*., 2021).

## Results

### Emergence with five years of nesting activity (five-year emergence)

Emergence of *M. tepidus timberlakei* at our perennial nesting site began on 17 May (2022), 25 May (2023), 29 May (2024), 10 June (2025), and 20 April (2026) (Fig. 4). Daily temperature (Fig. 5) and precipitation (Fig. 6) varied during the developmental periods for each of these five emergence years. Winter and spring heatwaves preceded the early 2026 emergence, with Tmax exceeding the 90th percentile of climate normals on 63 days during development. Precipitation was notably high in the winter. In 2022, which had the second-earliest emergence, late winter and spring heatwaves also occurred, although there were fewer exceptionally hot days (n = 43), and conditions were relatively dry. The middle emergence date, 25 May (2023), was preceded by cool, wet winter and spring conditions. The second-latest emergence, 29 May (2024), followed a cool, moderately wet spring, with some heatwaves in the previous fall and winter. The latest emergence, 10 June (2025), followed a very dry year with moderate temperatures, with only 16 days exceeding the temperature threshold, comprising two brief heatwaves in September and late February.

**Fig. 4–6.**
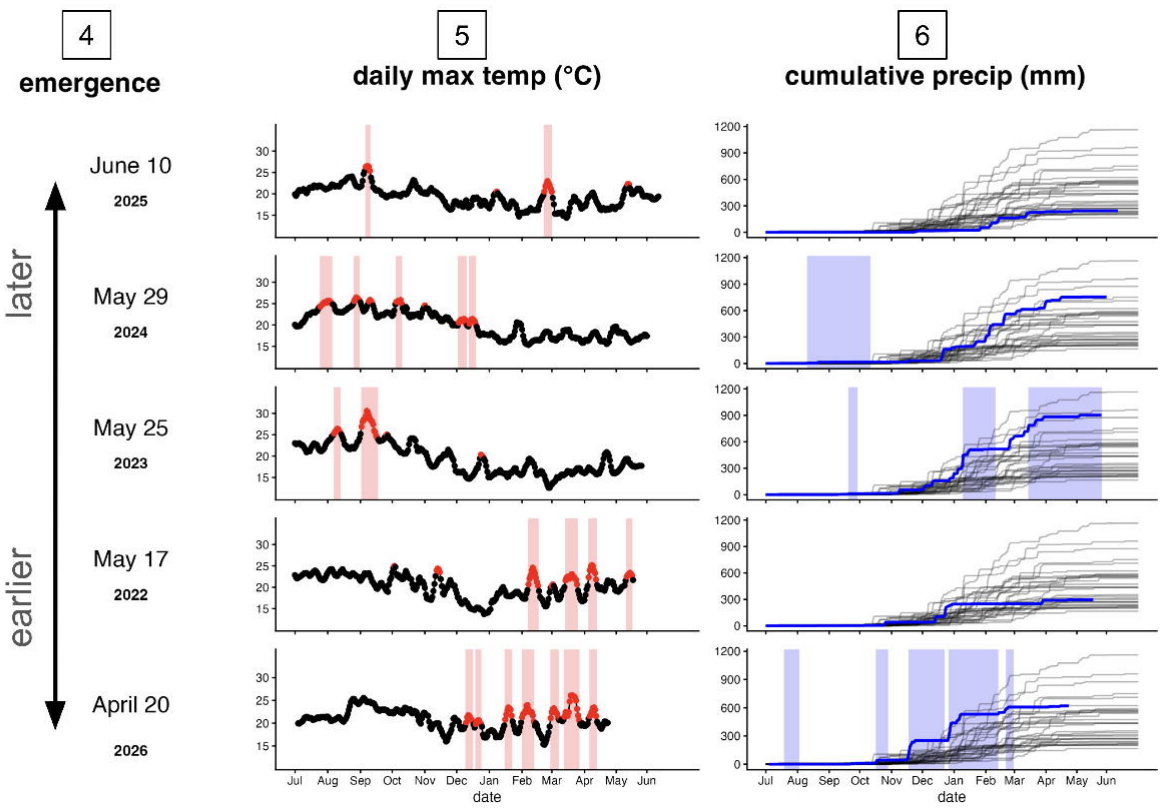
Emergence dates at a single nesting site in the previous five years range from April 20 in 2026 to June 10 in 2025 (**Fig. 4**). **Fig. 5** shows daily maximum temperature leading to emergence in each year as seven-day rolling averages. Red points indicate temperatures above the 90th percentile of the 30-year historical normals, meaning the plotted temperature exceeds 90% of historical values for that calendar day (1991–2021). Red shading shows time where temperature values were higher than 90% for five or more consecutive days. **Fig. 6** shows cumulative precipitation leading to emergence with blue lines. Thin gray lines are the cumulative precipitation values in the 30-year climate normals. Blue shading shows time when the observed cumulative precipitation values were higher than 90% of the reference normals for five or more consecutive days.

### Estimated emergence with 119 years of occurrence data (occurrence data)

We used 247 unique occurrence records from 1907–2026 as a proxy for *M. tepidus* emergence (Table S1). Estimated emergence dates ranged from 20 April (our 2026 observation) to 19 October, with a median of 13 July. We found a negative relationship between emergence timing and mean temperature, indicating that bees emerge earlier under warmer conditions (Fig. 7; LMM, β = –2.05 ± 0.60, t_85.7_ = –3.43, Bonferroni-adjusted p = 0.002, α = 0.05, R^2^_m_ = 0.06, R^2^_c_ = 0.14). The model estimates that emergence timing advances by approximately 2.05 days for every 1°C increase in mean temperature. The low R^2^_m_ value indicates that mean temperature alone explains relatively little variation in emergence timing. Nonetheless, the negative slope and low p-value indicate a significant association between temperature and emergence timing. In contrast, we found no relationship between emergence timing and cumulative precipitation (Fig. 8; LMM, β = 0.008 ± 0.01, t_187.6_ = 0.61, Bonferroni-adjusted p = 1.00, R^2^_m_ = 0.002, R^2^_c_ = 0.12).

**Fig. 7–8.**
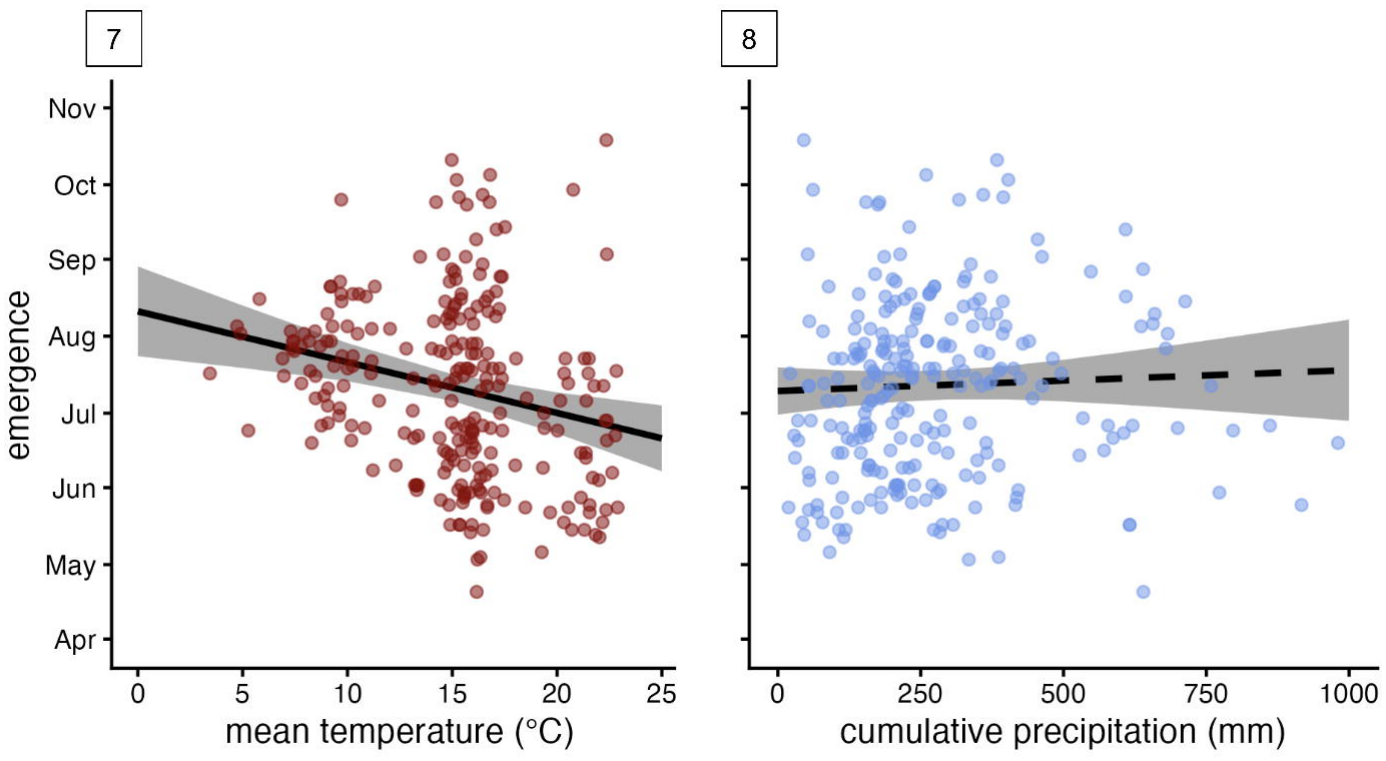
Bee emergence advances with temperature (**Fig. 7**), but not precipitation (**Fig. 8**) with 119 years of western United States occurrence data. In both panels, points represent *M. tepidus* emergence date proxied from occurrence data (n = 247) and climate (mean temperature or cumulative precipitation) during corresponding development time (July–May). Black lines and shading show LMM regression lines and 95% confidence intervals. Line type indicates relationship significance: a solid line represents the significant relationship between temperature and emergence (LMM, Bonferroni-adjusted p = 0.002), while a dashed line represents the insignificant relationship between precipitation and emergence (LMM, Bonferroni-adjusted p = 1.00).

## Discussion

Our 20 April 2026 observation of *M. tepidus timberlakei* was earlier than all previously recorded emergence dates at our perennial nesting site and in broader occurrence data across the western United States. In the previous five years at our nesting site, earlier emergence coincided with preceding heatwaves, whereas precipitation was more variable (five-year emergence). In occurrence data for *M. tepidus* across the western United States between 1907 and 2026, estimated emergence advanced with increasing temperature, albeit modestly, but had no relationship with precipitation (occurrence data). We included location in our models as a random rather than fixed effect to account for site-level variation in emergence timing while maintaining a focus on climatic emergence cues. Including additional fixed effects, such as geographic variables, could help explain additional variation in emergence timing.

Our temperature–emergence findings align with previous research (Table 1). However, this is the first study to examine this relationship in *Melissodes*, a large genus with 129 species distributed from Canada to Argentina (Wright *et al*., 2020). Understanding which environmental variables influence emergence is essential to predicting how phenology may change within this genus. This is especially relevant during global climate change (Seneviratne *et al*., 2021). In the western United States, climate conditions are shifting towards warmer temperatures, and early-season heatwaves, such as those preceding the early 2026 emergence, are becoming more likely (Jay *et al*., 2023). Together, these findings suggest that emergence may continue to advance and unusually early emergence events may become more common.

Because shifts in emergence can be deleterious when they lead to phenological mismatch (Freimuth *et al*., 2022), effective wild bee conservation requires anticipating these shifts through an understanding of environmental emergence cues. Anomalous phenological events, when documented and investigated, provide opportunities to build understanding of emergence cues. For example, a precipitation threshold associated with emergence in a spring-flying desert *Hesperapis* species (Melittidae) was identified after a novel fall flight period followed an uncharacteristically wet summer (Hurd, 1957). One limitation of inferring emergence cues from novel observations alone is that observed individuals may have immigrated or been transported from locations experiencing different weather conditions, as documented in dragonflies (Butler *et al*., 1975). Monitoring nest sites addresses this limitation by allowing researchers to quantify the environmental conditions experienced during development, rather than estimate them from the locations where flying adults are opportunistically collected. At nest sites, the use of emergence traps can ensure that sampled bees are emerging from the monitored nests rather than flying in from elsewhere (Hoge *et al*., 2026). Perennial nest sites can persist for several years or even decades in some solitary bee species (Danforth *et al*., 2019: Table 6-3). Thus, continued monitoring can inform emergence cues while accounting for site-specific variation.

Challenges remain in estimating historical emergence timing, as is also the case when estimating flowering onset in the better-studied field of plant phenology (Park *et al*., 2025). Our five-year emergence analyses address this challenge by using a monitored nesting site where emergence dates are directly observed, though this limits sample size. Our occurrence data analyses partly address this challenge by using only the earliest record of *M. tepidus* for each locality and year, because occurrence records may represent any point within the adult activity period, rather than onset alone. Using the earliest record for each locality and year provides a useful approximation of emergence timing, but this approximation becomes less reliable when sampling is limited. For example, our latest estimated emergence date (19 October) likely represents a specimen collected near the end of its flight window, rather than shortly after emergence. Although estimating emergence from occurrence data rather than a monitored nest site is a limitation, occurrence data allow for a much larger sample size than monitoring a single nesting site over multiple years. Novel methods for accurately estimating physiological age in preserved specimens could address this limitation by allowing emergence timing to be estimated from additional occurrence records, rather than relying exclusively on the earliest record for each locality and year. Wing wear is one such proxy for physiological age but becomes less accurate at higher wear scores (Mueller & Wolf-Mueller, 1993; O’Neill *et al*., 2015).

We found that emergence in a ground-nesting bee advanced with warming temperatures but was not associated with precipitation. Future studies incorporating additional species, specimens, and climate data will help determine how broadly this temperature–emergence relationship applies across bees. Additionally, we emphasize the importance of long-term monitoring at perennial nesting sites to identify emergence cues that cannot be distinguished using occurrence data alone. Because climate change will continue to increase the frequency and intensity of anomalous heat and precipitation events, incorporating knowledge of environmental emergence cues—such as temperature, demonstrated here—to predict phenological shifts in bees is essential for conserving bees, maintaining bee–plant synchrony, and ultimately sustaining pollination.

## Acknowledgements

Discussions of ground-nesting bee biology and nest site research with Bryan Danforth and other instructors and students at Bee Course 2026 bolstered and motivated this research. We thank students who monitored the ground-nesting site used in this study, including Liliana Lay, Juliana Goldman, Carys Goldsmith, Jorge De La Cruz, Everett Mathiason, and Victor Rodriguez. Jeremiah Bender photographed the *M. tepidus timberlakei* bee shown in Fig. 3. We acknowledge the time and expertise of all collectors, identifiers, and curators of bees used in this study, without whom this research would not have been possible. This work was supported by two National Science Foundation awards: 2139319 (Graduate Research Fellowship) to CNT and DBI-2102006 (Extending Anthophila research through image and trait digitization) to KCS.

## Supplemental Material

**Table S1**. Full dataset used in the occurrence data analyses (n = 247)

